# Accounting for heterogeneity in wild adult samples to measure insecticide resistance in *Anopheles* malaria vectors

**DOI:** 10.1101/2021.08.13.456216

**Authors:** Inga E. Holmdahl, Caroline O. Buckee, Lauren M. Childs

## Abstract

**Background:** Systematic, long-term, and spatially representative monitoring of insecticide resistance in mosquito populations is urgently needed to quantify its impact on malaria transmission, and to combat failing interventions when resistance emerges. Resistance assays on wild-caught adult mosquitoes (known as adult-capture) offer an alternative to the current protocols, and can be done cheaply, in a shorter time frame, and in the absence of an insectary. However, quantitative assessments of the performance of these assays relative to the gold standard, which involves rearing larvae in an insectary, are lacking.

**Methodology/Principal findings:** We developed a discrete-time deterministic mosquito lifecycle model to simulate insecticide resistance assays from adult-captured mosquito collection in a heterogeneous environment compared to the gold standard larval capture methods, and to quantify possible biases in the results. We incorporated non-lethal effects of insecticide exposure that have been demonstrated in laboratory experiments, spatial structure, and the impact of multiple exposure to insecticides and natural ageing on mosquito death rates during the assay. Using output from this model, we compared the results of these assays to true resistance as measured by the presence of the resistance allele. In simulated samples of 100 test mosquitoes, reflecting WHO-recommended sample sizes, we found that compared to adult-captured assays (MSE = 0.0059), larval-captured assays were a better measure of true resistance (MSE = 0.0018). Using a correction model, we were able to improve the accuracy of the adult-captured assay results (MSE = 0.0038). Bias in the adult-capture assays was dependent on the level of insecticide resistance rather than coverage of bed nets or spatial structure.

**Conclusions/Significance:** Using adult-captured mosquitoes for resistance assays has logistical advantages over the standard larval-capture collection, and may be a more accurate sample of the mosquito population. These results show that adult-captured assays can be improved using a simple mathematical approach and used to inform resistance monitoring programs.

**Author Summary:** Growing insecticide resistance in the mosquitoes that transmit malaria necessitates more widespread monitoring. Conducting assays on mosquitoes captured as adults is logistically simpler than raising them from eggs or larvae, the current recommended practice. However, this method is not widely used because survival when exposed to insecticide is known to depend on age and history of previous history as well as genetic resistance–factors that cannot be controlled when testing wild-caught adults. Here, we developed a mathematical model to quantify the difference in resistance measured via adult-capture assays compared to the gold standard larval-capture assays. We find that adult-capture assay results can be easily corrected using a formula based only on the measured resistance. This result has the potential to expand access to monitoring by reducing the time and infrastructure required to conduct these tests.

## Introduction

Insecticides used to target the *Anopheles* mosquito vectors of the *Plasmodium* parasite have long been a cornerstone of global malaria control. Most insecticides currently in use are pyrethroids, which are the only class of insecticide approved for safe use in bed nets (LLINs) and indoor spraying on walls (IRS)(1). The vast reduction of malaria transmission observed over the past few decades has been largely attributed to these interventions, but following increasing use of these tools–most intensely in sub-Saharan Africa–pyrethroid resistance is now widespread among malaria vectors(2). There is evidence that increasing resistance may be impacting the efficacy of these vector control tools, consistent with stalled progress in malaria control recently(3). This has been difficult to quantify, however, because of the difficulties of easily monitoring resistance in wild populations(4,5). Practical, cost-effective new tools for routine resistance monitoring are therefore urgently needed to quantify the impact of insecticide resistance on malaria transmission, and to combat failing interventions when resistance emerges.

Monitoring insecticide resistance is one of the main pillars of the Global Plan for Insecticide Resistance Management in malaria vectors (GPIRM)(6), and the WHO has published recommendations for prolonging insecticide efficacy, depending on local insecticide resistance levels. Resistance in *Anopheles* populations is currently categorized as a binary measure (resistant or not), based on a threshold of 10% of mosquitoes surviving exposure to insecticide after 24 hours, with sentinel sites monitoring populations at least annually. The current recommended protocol includes raising mosquitoes from collected larvae or to first generation (F1) progeny of captured adults, which necessitates access to insectary facilities and takes a week or longer. As a result of these requirements, in many malaria-endemic regions there are still large gaps in insecticide resistance monitoring data, both geographically and over time(7–9). Recent work has used spatial modelling to try and fill in these gaps across sub-Saharan Africa, by constructing annual maps of predicted resistance level by district, but many locations still have insufficient data, and geographic heterogeneity in resistance levels is likely to make inferred results difficult to interpret [7]. Furthermore, binary measures of resistance in particular make it difficult to understand trends over time and space, which is important in order to understand when and where control measures should incorporate new or alternate approaches, such as PBO nets(2,10).

Systematic, long-term, and spatially representative monitoring may require increased sampling of mosquito populations beyond the current sentinel site system. The WHO’s ‘Test procedures for insecticide resistance monitoring in malaria vector mosquitoes’ advises that control programs select sentinel sites that will be representative of both the eco-epidemiological zones and malaria intensities in their region(11). However, both of these variables are changing over time–as a response to malaria interventions, and because of climate change and changing land use(12–14). Thus, sentinel sites may not accurately represent insecticide resistance within a region. Furthermore, *Anopheles* populations appear to cluster at a relatively small geographic scale, due to reliance on freshwater breeding sites(6,15). For this reason, the GPIRM emphasizes the importance of monitoring and responding to insecticide resistance at the local level(6). Measuring more frequently, and with better geographical coverage, is therefore necessary in order to understand the extent and spread of insecticide resistance.

Resistance assays on wild-caught adult mosquitoes (known as adult-capture) offer an alternative to the current protocols, and can be done cheaply, in a shorter time frame, and in the absence of an insectary. Adult capture is not currently recommended by the WHO, however, due to heterogeneity of adult mosquitoes in age and exposure. The life history of a mosquito is known to impact its resistance to insecticides, and laboratory experiments have shown that insecticide resistance declines with age(16). Resistance also appears to decrease when mosquitoes have been exposed to insecticides before, weakening even resistant mosquitoes(17). Unlike mosquitoes raised from larvae, which are tested at a standardized age, mosquitoes captured as adults have unknown age at the time of testing and are likely to have a variety of insecticide exposure histories, so it is generally assumed that assay results would be difficult to interpret. Here, we examine the impact of heterogeneity in age and exposure history on the observed survival in insecticide resistance assays using a mosquito life cycle model of a mixed resistant and susceptible population, incorporating age and insecticide exposure-based mortality. We show how sampling from heterogeneous mosquito populations impacts insecticide assays taken using adult-captured mosquitoes compared to those using larval-captured mosquitoes, and evaluate both of these measures as predictors of true resistance in the population. We develop a simple correction that can be used as a guide to remove the bias in adult-captured assays, and propose that adult mosquito sampling offers a practical complementary approach for monitoring insecticide resistance in malaria-endemic regions.

## Results

Under our standard parameterization of the mosquito life cycle, we found that the mosquito population always reached equilibrium with the entire population either fully susceptible or fully resistant. Resistant equilibria were reached when insecticide coverage was high, and susceptible equilibria were reached when insecticide coverage was low (Fig 1A). The coverage threshold between susceptible and resistant equilibrium depended on the degree of spatial clustering. Higher spatial clustering led to a higher coverage required to reach a resistance equilibrium. When the spatial clustering parameter was equal to 1, such that the first blood feed exposure status determined all future exposures, the equilibrium population was fully resistant at 70% insecticide coverage or higher. When there was no spatial clustering, the population reached 100% resistance when insecticide coverage was at or above 30%. These trends in true resistance are reflected in the mean survival observed in the simulated resistance assays based on both larval- and adult-capture, although the adult-capture assays have lower survival at high resistance.

**Fig 1.**
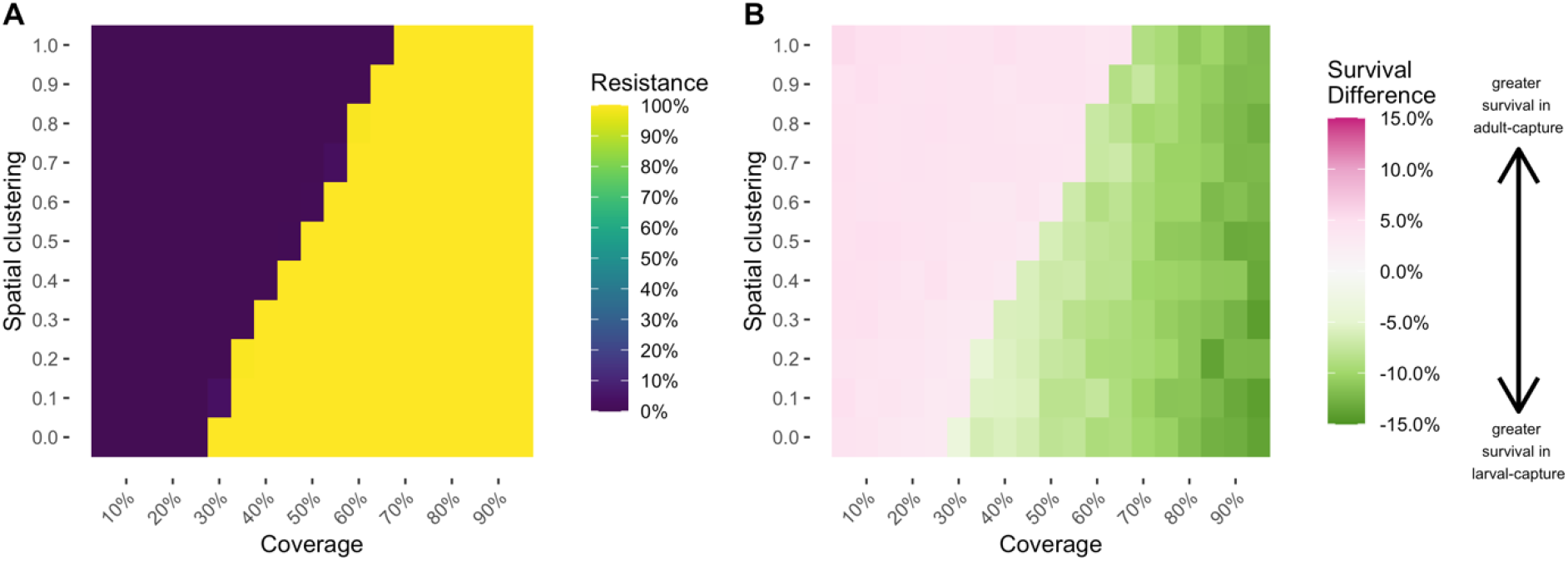
Difference between adult-capture and larval-capture assay results is determined by population resistance. A) Insecticide resistance across a range of insecticide coverage (x-axis) and spatial coefficient (y-axis) values at resistance equilibrium, 25 years following insecticide introduction. Color scale indicates % R alleles. B) difference between mean results from adult-capture and larval assays across a range of insecticide coverage (x-axis) and spatial coefficient (y-axis) values, conducted at the same time point. The color scale indicates the difference between mean assay results in adult capture compared to larval capture, i.e. adult capture survival – larval capture survival. Negative values indicate that observed survival is lower in adult capture than larval capture, on average; positive values indicate that observed survival is higher in adult capture than larval capture.

The survival difference between adult-capture and larval-capture methods maps directly onto equilibrium resistance values (Fig 1A and B), and does not appear to differ by spatial structure or coverage. We interpret this to mean that the bias in adult-capture assays depends on resistance proportion, and can be aggregated across spatial structure and coverage. When we remove grouping by these parameters, assays based on larval- or adult-capture both measured true resistance relatively well (Fig 2A). Both larval-capture and adult-capture assay results are well fit using linear models (larval model R^2^ = 0.9996, adult model R^2^ = 0.9960). Larval-capture assay survival is a very accurate measure of true resistance. Adult-capture assay survival is not a perfect correlate of true resistance: it overestimates resistance by 3-5% at low resistance (<15%) and underestimates by 10-12% at high resistance (>90%).

**Fig 2.**
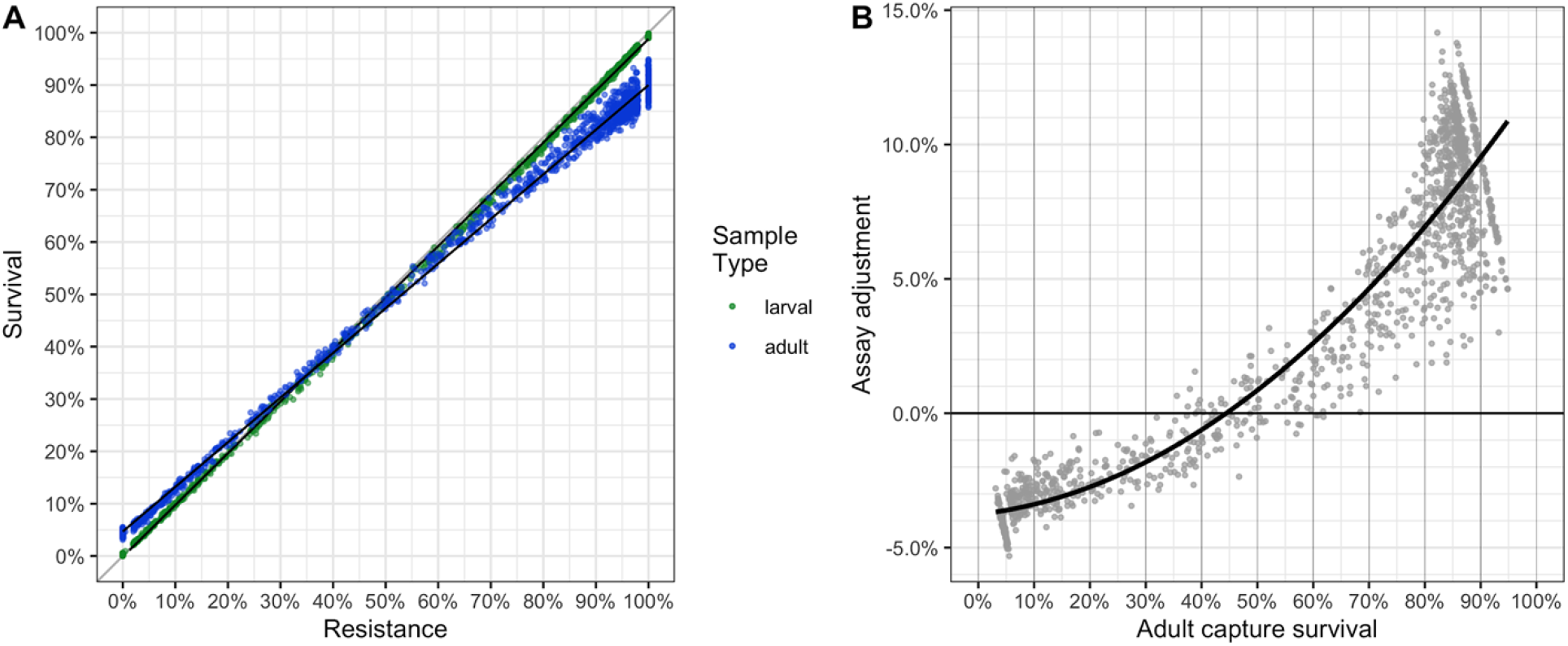
Resistance assay bias in adult-capture sample corrected with an adjustment model. A) Mean survival by true resistance, from 100 assays simulated with larval (green) and adult (blue) capture. B) Adjustment proposed for adult capture assays, to align with larval capture results which closely reflect true resistance frequency.

To construct an adjustment model, we subtracted the mean adult-capture results from the mean larval-capture results of each equilibrium sample (Fig 2B). We then modeled this survival difference against adult assay survival using a quadratic function (R^2^ = 0.9490).

The adjustment model was applied to adult-capture assays to improve the predictions of true resistance using observed adult survival. Using this adjustment (Fig 2B), the bias from adult-capture assays was reduced, only requiring information on the measured survival from those assays (Fig 2B). When we test the adjusted adult-capture results against the larval-capture results and against the original, unadjusted adult-capture results, the mean square error (MSE) of the adjusted adult-capture (0.0038) is lower than in the unadjusted adult-capture (0.0059) but is still greater than in the larval-capture (0.0018) (Table 2).

**Table 1.** Model performance comparisons.

|  | (Unadjusted) Adult-capture assay | Larval-capture assay | Adjusted adult-capture assay |
| --- | --- | --- | --- |
| Mean difference | -0.0132 | -0.0043 | -0.0073 |
| MSE | 0.0059 | 0.0018 | 0.0038 |

**Table 2.** Mortality parameters used in each assay simulation

|  | Genotype | Age- | Exposure | Insecticid |
| --- | --- | --- | --- | --- |

|  |  | based<br>mortality | history | e<br>mortality |
| --- | --- | --- | --- | --- |
| Adult-capture: controls | X | X | X |  |
| Adult-capture: exposed | X | X | X | X |
| Larval-capture: controls | X | day 3 only |  |  |
| Larval-capture: exposed | X | day 3 only |  | X |

### Sensitivity analyses

In sensitivity analyses, testing our assumptions of the mosquito life cycle and effects of insecticide exposure, larval capture remained a very good predictor of true resistance (Figs S1-6). Under all sensitivity analysis the relationship between adult-capture and larval-capture assays followed a similar trend as shown above, with small variations. The models with low larvicide exposure, a lower (intermediate) mortality related fitness cost, and no effect of insecticides on longevity garnered nearly identical correction curves (Fig 3).

**Fig 3.**
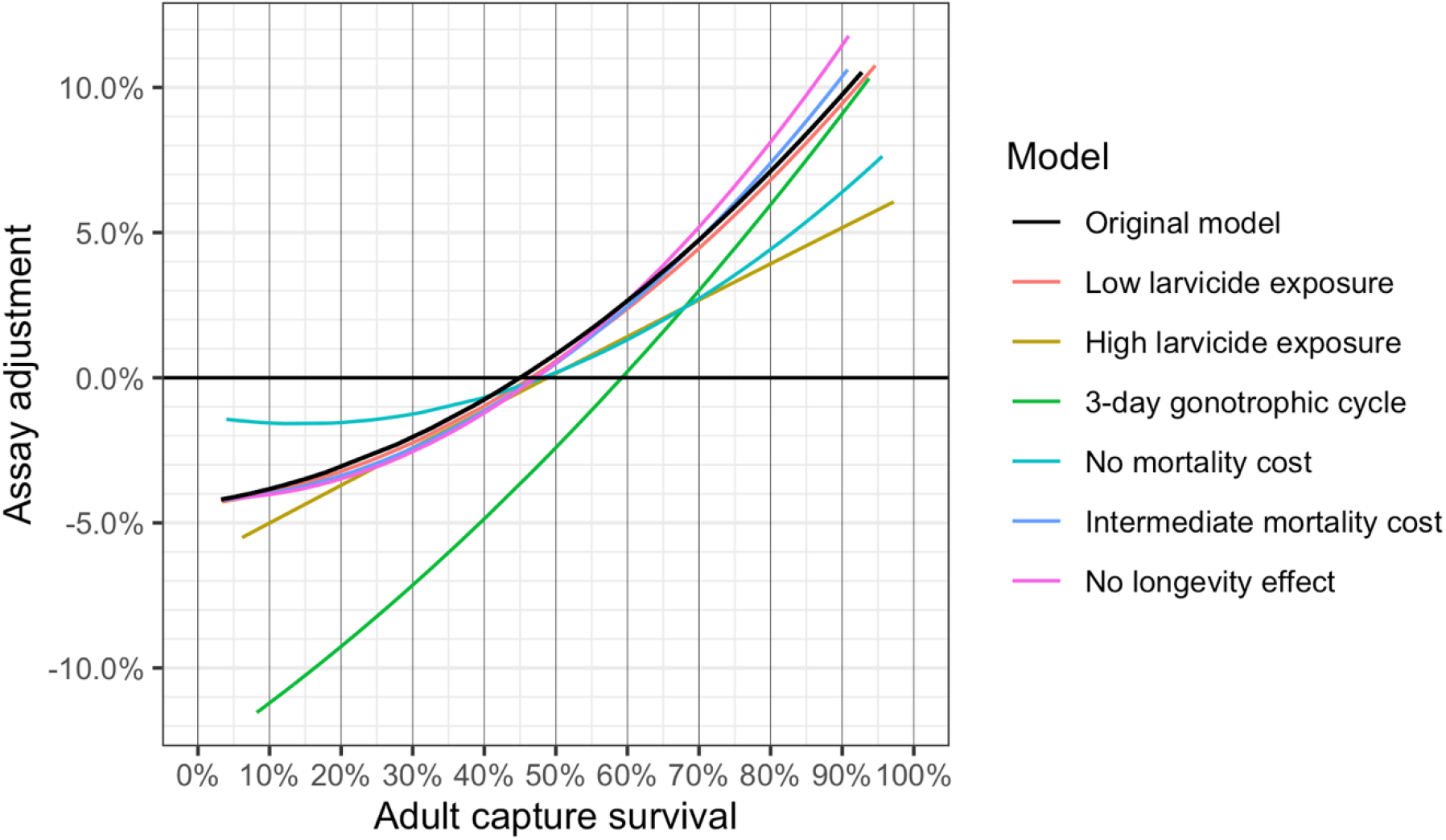
Adjustment model follows a similar form across sensitivity analyses. Adjustment curves generated from each of the models run in the sensitivity analyses, plotted against the results of the original model (in black). Within each sensitivity analysis, the above curve shows the difference between the adult-capture assay results and the larval-capture assay results, which is a good measure of true resistance.

The mean resistance measured using adult-captured assays were always higher than larval-capture assays at low resistance, and lower than larval-capture assays at high resistance (Fig 3). The resistance level at which adult-capture assays went from being an overestimate to an underestimate compared to larval-capture assays–i.e., where the two mean values were equal – ranged from 46.5%-49% resistance for all analyses apart from shortened gonotrophic cycle, which intersected at 59.5%. Adjustment curves generated in sensitivity analyses diverged from those in the original analysis the most at very high or very low resistance levels, in particular, for those with high larvicide exposure, 3-day gonotrophic cycle, and no mortality related fitness cost. At higher resistance levels, mean adult-capture survival was up to 11% lower than mean larval-capture survival, in the model with no effect of insecticide exposure on longevity (Fig 3, pink line). At lower resistance levels, the mean adult survival was up to 11.5% higher than the mean larval survival, in the model with a 3-day (rather than 4-day) gonotrophic cycle. The adjustment curve generated by the model with a shorter gonotrophic cycle had the greatest divergence from the other model results, leading to a much larger overestimate of resistance in the adult-capture assays at low levels of resistance (Fig 3, green line).

## Discussion

Insecticide resistance represents a major threat to the gains in malaria control achieved over the last several decades. Monitoring insecticide resistance over time and in different settings is an important component of surveillance for malaria control programs, and current approaches are lacking in this regard, partly because of the impracticality of current resistance assay protocols. We used a mathematical model of a mosquito population to compare the results of simulated adult- and larval-capture assays, and found that the bias in assays using adult-capture can be reduced using a simple correction based on measured resistance alone, allowing us to align adult-capture assay results more closely with larval-capture assay results. After adjusting for this bias, adult-capture can be a consistent measure of overall resistance–an approach that may better inform resistance monitoring efforts.

Adult-capture is likely a much more practical strategy for insecticide resistance monitoring than the larval-capture assays, which is the current recommendation by the WHO. Adult-capture can be done without relying on resources such as insectaries, and is therefore more portable, allowing for wide scale testing in more remote and varied locations. In addition, they take much less time–tests can be conducted immediately after collection, as opposed to the week or more required to raise larvae or F1 progeny to the correct age in the insectary. These advantages could allow for a shift away from the sentinel site model, towards more systematic monitoring over geographic regions.

In addition to logistical advantages, adult-capture may also be more representative of the mosquitoes that are contributing to malaria transmission (6). In theory, as we see in our model, larval-capture results are a very good measure of true resistance. However, this depends upon the assumption that they are drawn from the same population that we are interested in, i.e. the mosquitoes that are likely contributing to malaria transmission. This may not be the case in the field: larval collections, in particular, may overrepresent very few mosquitoes (due to egg laying behavior) and adult-capture can more easily target mosquitoes that are actively seeking out humans for blood feeding.

This model makes several simplifying assumptions, and relies on a single data set to parameterize the mortality of resistant mosquitoes following multiple exposures. We do not model multiple *Anopheles* species, which often overlap in a single ecological setting. It is possible that unknown characteristics of these or other traits of field-captured adults may lead to a different relationship between resistance and adult-capture survival than we observe in our model. Some field experiments have demonstrated higher resistance in adult-captured mosquitoes than measured in the birth cohorts, which appears counter to the results of the lab work in Viana et al.(18). However, these data may not be in conflict: in our model, we show that, depending on true population resistance, adult-capture can yield both higher and lower mortality than larval-capture. However, it is possible that we are not fully discerning the fitness landscape of insecticide resistance. While there are currently very little data available to compare this in the field, experiments pairing larval- and adult-captured assays in regions with different levels of true resistance could lend support to either of these hypotheses.

The results of this model demonstrate that adult-capture, through a simple bias correction, can be a useful and consistent measure of resistance. Field experiments should be conducted to validate our modeling results, allowing adult-capture to be prioritized moving forward as a more practical strategy for wide scale testing than the methods currently recommended.

## Methods

### Mosquito life cycle model

We developed a discrete-time deterministic mosquito lifecycle model that includes a single resistance gene and allows for multiple exposures to insecticide. Insecticide exposure is assumed to occur only through LLINs, and therefore only on blood feeding days which occur on every fourth day (labelled “feed” in Figs 4 and S7). Resistance status is determined at a single locus, by *S* and *R* alleles. We assume three phenotypes: full susceptibility (*SS*), intermediate resistance (*SR*), and resistant (*RR*). In this population, the phenotypic characteristics of the SR heterozygous mosquitoes are exactly intermediate between the SS and RR homozygous phenotypes. Offspring genetics follows Mendelian inheritance patterns, with random mating.

**Fig 4.**
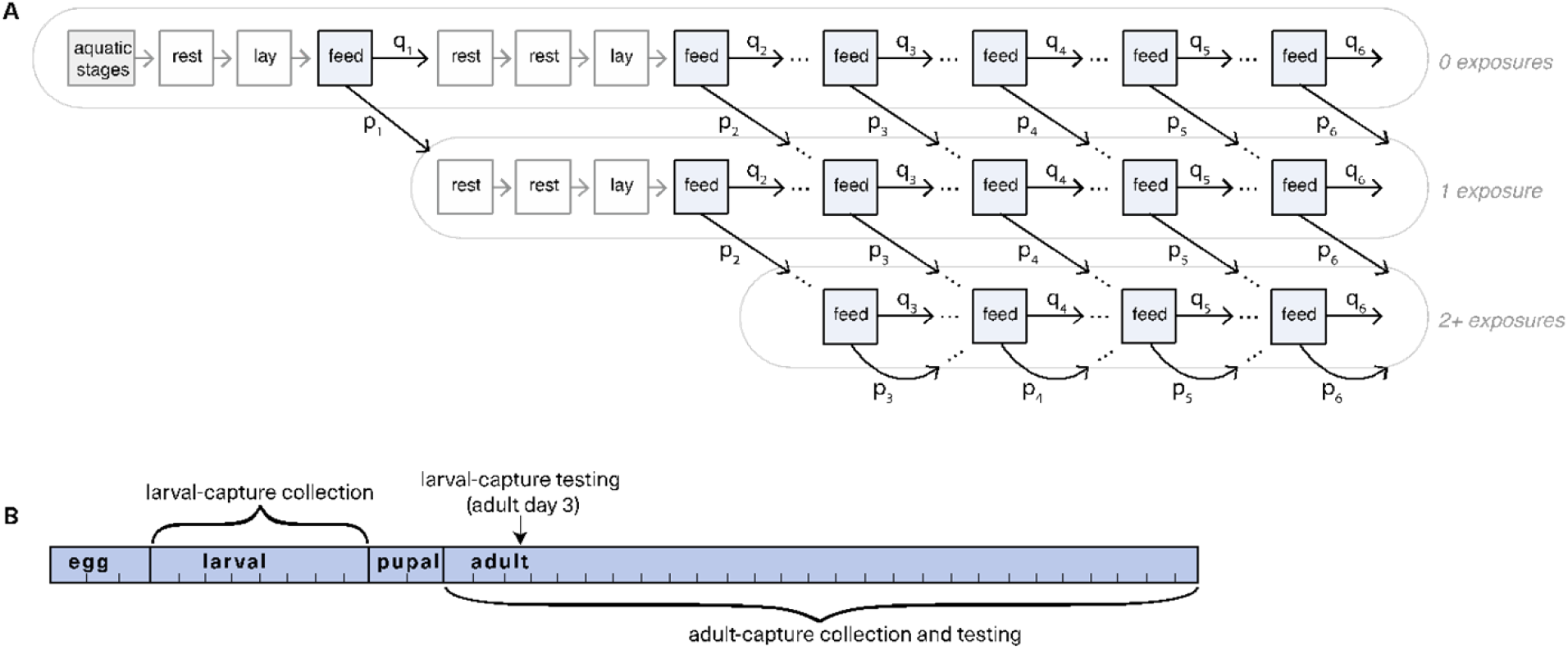
Mosquito life cycle model and sampling schematics. A) Simplified schematic of mosquito population model for a single genotype, with exposure days highlighted blue. The aquatic stage (in grey) is split into three components: the egg stage is 3 days, larval stage is 5, and pupal stage is 2 days. In adult (non-aquatic) stages, each compartment represents a single day. Mosquitoes have the opportunity for exposure on each feeding (highlighted), which varies by spatial coefficient s and is determined by past exposure history. The number of past exposures is not tracked beyond 2 exposures. Mosquitoes live up to 26 days. The first full gonotrophic cycle is shown, while later cycles are abbreviated to show only feeding days (i.e. rest and laying days hidden). The fully elaborated diagram is shown in Fig S1. Daily mortality and egg laying also not shown for ease of visualization. B) Timeline for larval-capture (upper) and adult-capture (lower) collection and testing.

We modeled the mosquito life cycle in single day timesteps: mosquitoes lay eggs every fourth day, three days after blood feeding (Figs 4 and S7). In every stage of the life cycle, a proportion of mosquitoes die each day. In egg and pupal stages, mortality is constant (Table S1); in the larval stage, it is density dependent, following a Ricker-type nonlinear survival function (19). In the adult stage, mortality is based on age, genotype, and exposure history (Fig S8). In addition, there is immediate mortality on feeding days, induced by insecticide exposure due to encountering an LLIN. We incorporated effects of past insecticide exposure on mortality, as documented in lab experiments in (17), through two mechanisms: increased immediate mortality upon exposure to insecticides, and an increase in daily age-based mortality for the rest of the mosquito’s life following exposure. The homozygous resistance (RR) mortality phenotype is from the resistant TIA strain data in (17). The homozygous susceptible (SS) mortality phenotype is based on an entirely susceptible lab strain from (17,20). Gompertz-Makeham survival curves were fit to these mortality data (see Supplemental Methods) to obtain both immediate and daily mortality estimates for mosquitoes with a history of insecticide exposure and without (Table S2 and S3). The differences between these mortality data impose a fitness cost of resistance.

### Insecticide exposure

Mosquitoes in this model blood feed up to six times, so have a maximum of six insecticide exposures. However, to simplify the model, we only track up to two past exposures explicitly. Insecticide coverage is a parameter input for each simulation, and is used to determine the probability *p(f)* of insecticide exposure at each blood feed in the absence of heterogeneity, where f refers to the blood feed count. To account for outdoor biting and net failure in the probability of exposure, we scale coverage by “LLIN effectiveness” (80%). The probability of exposure on the first feed p(1) is always equal to the coverage multiplied by the LLIN effectiveness.

To capture spatial heterogeneity and avoid unrealistic coverage of LLINs (see Supplemental Methods), we introduced a proxy spatial structure(15), which causes the probability of future insecticide exposures to depend on a mosquito’s past exposures. This reflects that mosquitoes may tend to feed locally rather than biting the human population randomly. To do this, we used a spatial clustering parameter *s* to modify the probability of exposure on all blood feeds *f* after the first feed (Fig 4). We varied *s* between 0 and 1: when *s* = *0*, there is completely random mixing in the model and the probability of exposure on each feed is constant across all feeds (i.e. always the same as in the first feed). When *s* = *1*, there is perfect spatial clustering and exposure on future feeds is determined entirely by the exposure status on the initial feed.

Intermediate exposure probabilities are calculated as follows, depending on previous exposure:

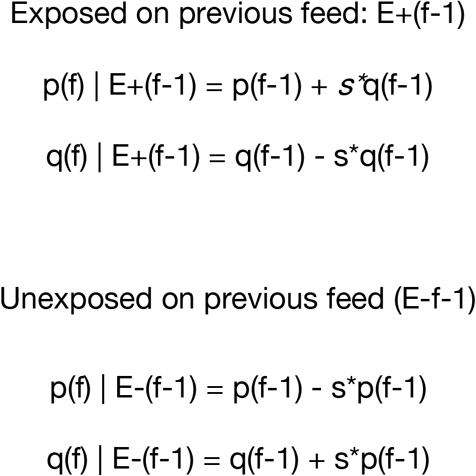

For some blood feeds there is more than one way to compute the probability of exposure (e.g., the third blood feed by a mosquito that had a single previous exposure could have been either exposed or unexposed on the most recent blood feed). In these cases, we computed all of the possible exposure probabilities and used their mean.

Both spatial coverage and the proportion resistance in the population impact how coverage maps onto the actual proportion of blood feeding mosquitoes exposed on a given day (Fig S9). When insecticide resistance is low, and there is at least some level of spatial clustering (s>0), a disproportionate number of the mosquitoes alive at a given time will be those that were not exposed in their first feed because all of the susceptible mosquitoes that were exposed on the first feed died. This leads the proportion exposed to be much lower than suggested by the coverage and LLIN effectiveness (Fig S9A). However, when the population is highly resistant, few mosquitoes die on their first or second blood feed, and the proportion of feeding adults exposed on a given day match more closely to the product of coverage and LLIN effectiveness (Fig S9B).

### Insecticide resistance assay simulations

As the model is deterministic, under each spatial coefficient *s* and insecticide coverage, the population reaches a single equilibrium proportion of the population that is resistant. True resistance in the modeled population is calculated as the resistance allele frequency in the whole population:

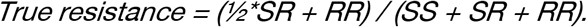

To compare insecticide susceptibility assay results under different collection strategies, we simulated assays by drawing individuals from the constructed population. To conduct an individual assay, 200 mosquitoes were sampled randomly from the population. 100 of these mosquitoes were exposed and 100 were used as controls. The probability of mortality for each mosquito sampled for the assay is a function of age, exposure history, genotype, and whether they are in the exposure or control sample (Table 1). To determine whether an individual mosquito dies, we drew from a binomial distribution with that probability. For adult-captured assays, we drew mosquitoes randomly from the adult population on the day of collection to determine their genotype, age, and exposure history. For the larval-captured assays, we drew mosquitoes randomly from the larval population on the day of collection to determine their genotype, and then assumed that in the assay they have the mortality probability of a three day old adult. This would be equivalent to collecting as larvae and raising them in an insectary, and performing the assay three days after emergence.

To adjust for mortality observed in the control, the results of each assay were corrected using Abbott’s formula (21):

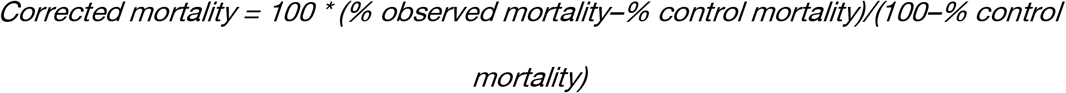

While the WHO only recommends using Abbott’s formula when control mortality is > 5%, we used the adjustment regardless of control mortality in order to directly compare adult to larval sampling. We were interested in corrected survival, which is 1-corrected mortality.

### Comparison of sampling methods

To compare adult-capture to larval-capture, we ran the model starting with 1% resistance in the population until it reached resistance equilibrium, across each value of insecticide coverage and spatial coefficient. For any combination of these parameters, resistance at equilibrium was either 0% or 100%. To evaluate the differences in collection strategies at intermediate resistance, we sampled from the mosquito population over time. To exclude the instability that occurs early in simulations with high coverage and ensure that we were using mosquito populations that, while not at equilibrium, were at least changing slowly, we excluded the first year of each simulation. From the remaining data, we created a sampling dataset, by choosing 10 days randomly for each combination of coverage and spatial clustering parameter from which to draw samples.

For each day that samples were drawn, we simulated adult- and larval-capture resistance assays 100 times each. We took the mean of the corrected survival across these 100 samples. We are interested in the difference between the results from the adult-capture and larval-capture assays, so we calculated the “survival difference” by subtracting the mean survival using adult-capture from the mean survival using larval-capture. By this measure, a positive survival difference value means that the observed mean survival is higher in adult-capture assays than larval-capture assay, and a negative survival difference means that survival is higher in larval-capture assays.

### Adjustment model

We compared the results of each of the mean larval- and adult-capture assays with the true resistance proportion, and used linear regression to model these relationships. We then built an “adjustment model” by using linear regression to model the relationship between survival difference and the results of adult-capture insecticide assays. This “adjustment model” takes adult survival as an input and returns the difference between adult-capture survival and larval-capture survival. The value of the difference is added to the adult survival to create the *adjusted adult-capture assay result*, our new estimate of resistance.

To evaluate the efficacy of this adjustment model, we use the sampling dataset described above. We then re-ran our insecticide assay simulations, conducting 100 samples from each of these scenarios. We used the adjustment model on the adult-capture assay results to calculate the adjusted adult-capture results. To compare the accuracy of this measure against the accuracy of the larval-capture and unadjusted adult-capture assays, we calculated the mean square error (MSE).

### Sensitivity analyses

In this model, we made important simplifying assumptions, including that insecticide exposure only occurs via LLINs (as opposed to indoor residual spraying, larvicide, and agricultural exposure). In addition, resistance in this model is determined at a single, two-allele locus for simplicity, whereas insecticide resistance may have multiple causes of varying genetic complexity(22–25).

We conducted several sensitivity analyses to examine assumptions of our model around mosquito life cycle parameters and the effects of insecticides. We examined our assumption that there is a fitness cost due to earlier mortality in RR and SR compared to SS mosquitoes by repeating our simulations using the fully resistant (RR) mortality curves for all genotypes in the model, and again using an intermediate mortality curve for the SS mosquitoes–by using the mean of the RR and SS mortality for each day (Figs S1 and S2). We also looked at the sensitivity of our results to the assumption that there is a longevity effect (increasing daily mortality) in resistant mosquitoes that survive exposure. To do so, we assumed the only effect of past exposure was to increase immediate mortality on the day of exposure (Fig S3). In addition, we examined our assumption that all mosquitoes have a strict 4-day gonotrophic cycle, by instead using a 3-day gonotrophic cycle (Fig S4). While neither of these perfectly represent the blood-feed frequencies in the field, which vary, this allows us to understand the impact of this assumption on our model results. In both scenarios, mosquitoes are capped at six gonotrophic cycles, and the maximum lifespan is 26 days. Finally, we ran two sensitivity analyses where we incorporated insecticide selection during the larval stage, to reflect the possibility of environmental larvicides, imposing either a low or high additional mortality rate on each day of the larval stage (Figs S5 and S6).

